# Integrated Cell Expression Toolkit for low input transcript profiling

**DOI:** 10.1101/458851

**Authors:** Robert C. Day, Tanis D. Godwin, Chris Harris, Peter Stockwell, Anthony Shaw, Parry J. Guilford

## Abstract

Several methods exist for constructing sequencing libraries from single-cell amounts of RNA. However, different library preparation methods are required to enable full length or strand specific 3’ end profiling. The Integrated Cell Expression Toolkit (ICE-T) maximizes functionality by integrating these approaches with novel strategies that provide a range of profiling options.

Transcriptome profiling at high cellular resolution has many applications in biology but is hampered by a need to boost the representation of RNA to guard against losses during processing and enable efficient use of high throughput technologies. RNA amplification techniques developed for profiling limiting amounts of RNA by microarray have recently been adapted to NGS^1-5^. These various, but distinct, techniques enable both full length profiling^3^ and strand specific 3’ end profiling^2, 4^. Here, we improve flexibility and capability of low input RNA sequencing by consolidating library preparation protocols and exploiting synergies obtained by using different library preparation methods on the same sequencing run.

The ICE toolkit uses a cDNA synthesis primer that enables RNA amplification by either *in vitro* transcription (IVT) or PCR (Supplementary Sheet 1). Smart Seq2^3^ (SS2_PCR) and CelSeq^2^ (CS_IVT) were selected as the component core methodologies due to their ability to enable high sensitivity with full length sequencing and inexpensive multiplexing of strand specific 3’ end^1^ libraries respectively. Unlike IVT based protocols, a single SS2_PCR reaction is able to generate useful amounts of amplified product from single cell amounts of total RNA. SS2_PCR however relies on a double transposition that removes the ends of the library including any inline barcodes added during cDNA synthesis. This limitation led to the development of our PCR_IVT approach which maintains the sensitivity and input flexibility of PCR whilst maintaining inline barcoding.

ICE-T therefore enables three core options for library preparation: CS_IVT and PCR_IVT both generate stranded data limited to the 3’ end of transcripts and SS2_PCR generates non-stranded libraries conducive to full length profiling (Fig. 1 and Supplementary Fig. 1). Aspects of the pre PCR cDNA synthesis reaction such as template switching and buffer composition were also explored (Supplementary Fig. 2 and Supplementary Note).

**Figure 1.**
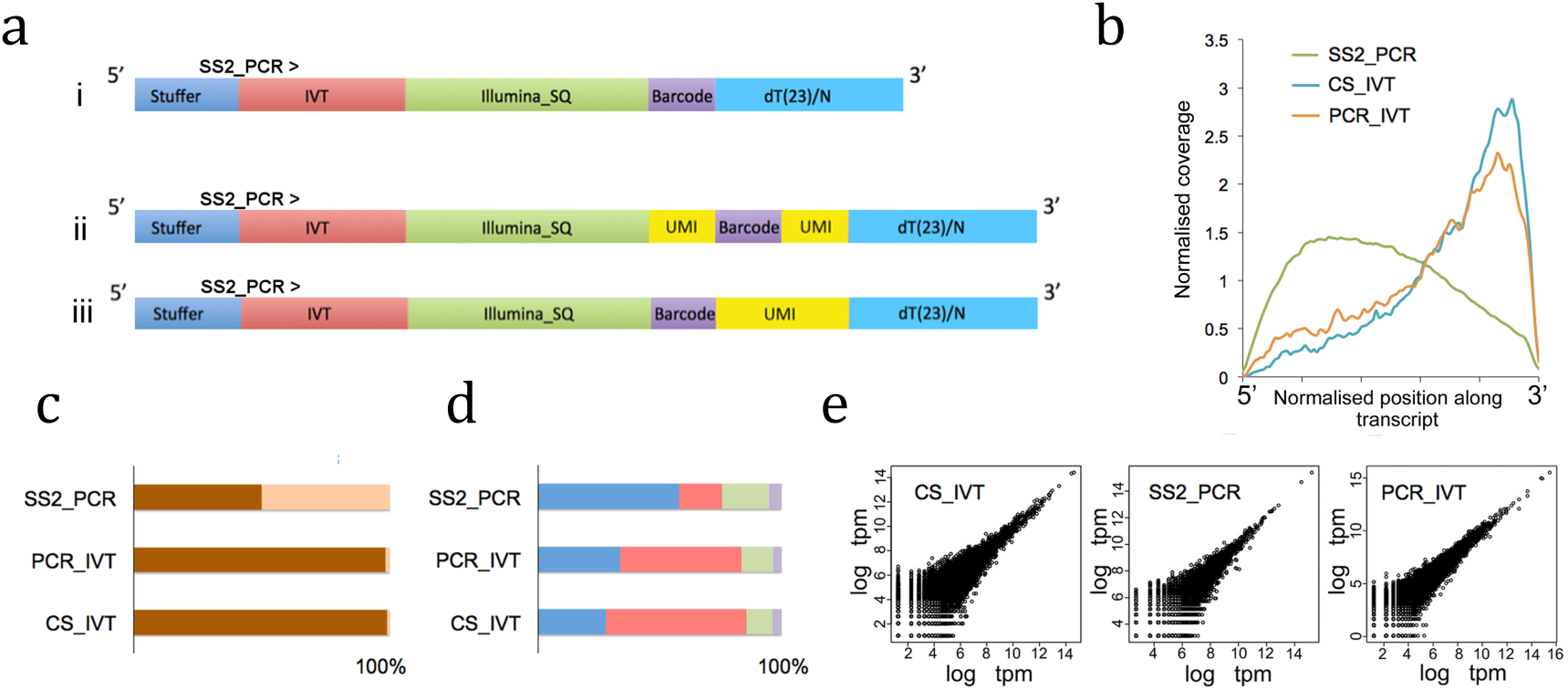
Comparison of ICE-T RNA sequencing strategies. A, i The basic cDNA synthesis primer design enables both PCR and/or IVT based amplification. Inclusion of a string of random bases, as in ii and iii, helps combat low complexity and enables molecular counting by universal molecular identifier (UMI). B, Different library types generate reads with mapping biases towards different regions of the transcript. They also vary by the percentage of reads mapping to each strand, C: Dark brown and Light brown denote sense and anti-sense strands, respectively) by the percentage of reads mapping to different genomic features, D: Blue, red, green and purple denote mappings to exonic, UTR, intronic and intergenic regions, respectively). E: Replicate data for each strategy tested showed a high degree of reproducibility (log tpm=log base 2 transcripts per million and input ~1 ng RNA).

To illustrate the flexibility and utility of ICE-T, we designed an experiment that integrated several workflows within the ICE toolkit (Fig. 2). The sequencing run combined PCR_IVT and SS2_PCR libraries to profile the transcriptomes of single cells and cell mixtures. 40 individual samples were sequenced as 3’ end libraries that contained ERCC spike-ins to help assess sensitivity and amplification biases of the PCR_IVT protocol. Analysis of the ERCC^6^ spike-in showed counts generated by PCR_IVT were directly proportional to the concentration of input RNA; approximately 48% of input RNA molecules were represented in our libraries (Supplementary Fig. 3), a representation similar to that seen in recent studies^2, 3, 7-10^. In addition, PCR preamplifications for each sample type were combined and made into representative full length Nextera libraries using the SS2_PCR approach.

**Figure 2.**
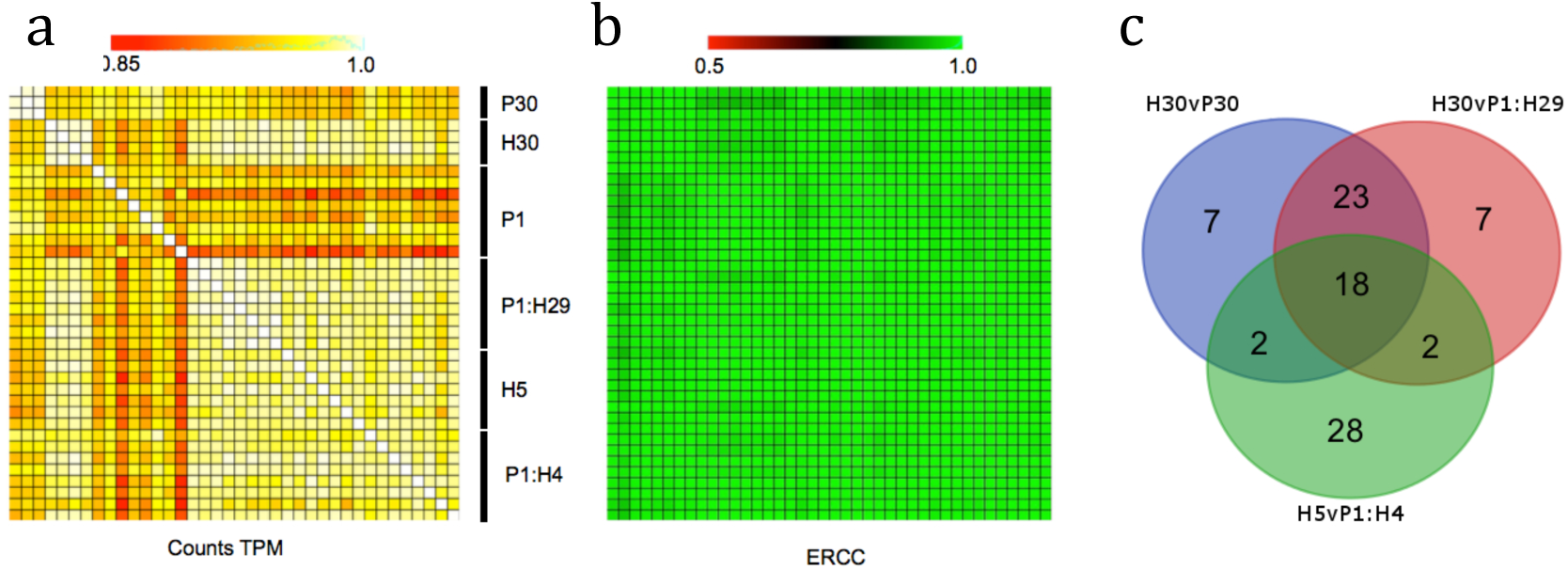
PC3 and HeLa cell mixture experiment. A, Correlation analysis of the count data (tpm=transcripts per million) mapping to genes or B, the ERCC spike in. P and H denote PC3 and cells respectively. C, Tracking of the top 50 most differentially expressed genes in the PC3 direction (based on fold change and p_value < 0.005) when comparing 30 PC3 to 30 HeLa cells (Blue) tracked in cell mixtures containing one PC3 cell with four HeLa (Green) and one PC3 cell with twenty nine HeLa (Red).

The use of individual PCR preamplifications enabled us to assess each sample by Bioanalyser trace (Supplementary Fig. 2 B,C) and PCR yield. Aliquots of individual PCR preamplifications can be stored for further downstream investigations, but they also provide an opportunity to adjust sample representation. In the experiment presented in Figure 2, we doubled the IVT input for our single cell samples to boost the number of reads relative to the other samples (Supplementary Fig. 4). Estimates for the number of RNA molecules per cell obtained from single cell PCR_IVT data from PC3 and LNCaP cell lines were in accord with similar studies^2, 7, 8^ (Supplementary Fig. 5) and delimitation between different cell types was straightforward^11^ (Supplementary Fig.6 and Supplementary Sheet 3).

We envisage one application of ICE-T to be the detection of minority cells in mixed cell populations. To do this effectively robust markers are required with strong differential expression between component cell types and with expression that is largely resistant to biological and technical variability. To explore this concept, we compared transcriptomes from samples of thirty PC3 cells (P30) and thirty HeLa cells (H30) to identify genes with strong differential expression (Supplementary Sheet 3). We then tracked the fifty most differentially expressed genes in the PC3 direction in mixtures containing a single PC3 cell mixed with either four (expected to show high levels of technical and biological variation) or 29 HeLa cells (expected to have low variation but a high dilution of the minority cell). These are termed P1:H4 and P1:H29 respectively (Fig. 2 C and Supplementary Sheet 3).

Only 40% of the 50 genes were still in the top 50 differentially expressed list for the P1:H4 samples whereas 82% were still in the top 50 in the P1:H29 samples. Using a p-value cut off of p=0.001, these numbers drop to 12% and 80% respectively. The results indicated that despite a higher dilution of the minority cell type in the P1:H29 samples minority cell screening is more robust using 30 cell aliquots than using 5 cell aliquots. Furthermore, six genes correctly indicating the presence of the minority cell in both dilution mixtures (*UCHL1*, *PRSS3*, *MAGED1*, *HIPK2*, *DUSP6* and *ARHGAP29*) effectively model the discovery of single gene markers for PC3 that are largely immune to technical and biological variation in our system.

In some instances, it is necessary to follow up low pass sequencing with increased sequencing depth of a few select samples. This usually involves inline barcodes that have been added prior to knowing which samples are to be combined for the subsequent low complexity run. Using a low number of samples can reduce the diversity during the barcode read and overlapping clusters may not be separated by the software. Highly biased sequence composition may also have deleterious effects on the signal intensity calibration reducing quality metrics. Conventional approaches to alleviate such problems is to dilute the library to the point where very few overlapping clusters are found and/or by adding a PhiX control library (reducing the effective concentration of the biased library, and introducing some added diversity). However both approaches reduce the yield of useful data.

ICE-T uses two different strategies to maximize biological information obtained from a sequencing run whilst helping to ensure complexity of the inline barcode position. Our preferred strategy is that of a blended run that involves a mixture of 3’ end and full length libraries on the same MiSeq run as already described. Since the Nextera tagmentation process inserts sequencing linkers into random sites across the DNA, reads initiate from random positions. Addition of these libraries to 3’ end libraries is therefore largely equivalent to adding a PhiX control i.e. adding complexity to Read1 in positions adjacent to the inline barcodes and dT tail of the primer. However, if SS2_PCR libraries are not part of the experimental design, we provide an alternative cDNA primer design that can be used alongside the standard cDNA synthesis primers. These incorporate random bases adjacent to the standard inline barcodes to alleviate complexity issues (Fig. 1A).

The random bases in the optional cDNA synthesis primer also act as unique molecular identifiers^7, 12^. Thus, use of these primers enable amplification biases to be assessed via molecular counting. Molecular counting from our own low pass experiments indicated that most reads are derived from discrete RNA molecules. Molecular counting may therefore be largely redundant if multiplexing many samples on a single MiSeq run (Supplementary Fig. 7).

Low pass sequencing is highly efficient with regard to gene detection (Supplementary Fig. 8) and identification of different developmental programs^9^. However, the investigation of hundreds of cells on a MiSeq run requires additional barcoding. A recently described a Tn5 transposase system uses the combination of inline barcodes and a transposase added barcode for use on the Fluidigm C1 system^7^. The authors created a library of XTn5 loaded with different barcoded linkers enabling creation of dual barcoded libraries suitable for 3’ end sequencing. However, creating large preparations of Tn5 and large numbers of aliquots loaded with different barcodes isn’t a feasible approach in many laboratories. Here we describe an alternative Tn5 based approach using a commercially available Tn5^13^ (Creative Biogene, NY). We loaded the Tn5 with annealed oligo preparations conducive to adding a standard Illumina i7 linker and added barcoded Nextera i7 adapters in the subsequent library finishing (Supplementary Fig. 9). Libraries barcoded in this way required an additional adapter trimming step during data processing that removed an ~5-10% of the raw reads (data not shown), however the generated data was highly reproducible and facilitates the use of hundreds of samples on a single MiSeq run.

ICE toolkits use of accessible reagents and well established molecular techniques will enable rapid adoption of this methodology. Component reactions of the ICE toolkit are easily transferrable to ultra-high throughput pipelines via miniaturisation and automation. However, the current implementation is focused on low to high throughput profiling experiments. ICE-T is particularly well suited to generating sensitive transcriptome maps using topographical RNAseq^14^ and an integrated deployment of its different library styles presents a powerful solution for experimental systems where no reference genome is available; essentially generating a reference transcriptome and efficent transcript counting in a single run.

## METHODS

Methods and any associated references are available in the online version of the paper Accession codes. Raw sequence reads are available at the NCBI SRA database (SRP067618).

## ACKNOWLEDGEMENTS

The authors wish to thank Mik Black and Tyler McInnes for help processing count data using R. Les McNoe and Caroline Beck for constructive comments on the manuscript. This work was supported by an MBIE Smart Ideas grant to PJG.

## AUTHOR CONTRIBUTIONS

RCD designed and tested the toolkit with help from TDG and CH. RCD and PS developed the data processing pipeline. RCD carried out differential expression analysis and made the majority of the figures. TDG harvested cells and maintained tissue cultures with CH. CH carried out PCA analysis and generated the corresponding figures. AS did the ERCC analysis and generated the appropriate figures. PJG helped coordinate the study and assisted RCD to write the manuscript.

## Cell culture

LNCaP, PC3 and HeLa cancer cells were obtained from ATCC. All cell lines were maintained as adherent cultures. The LNCaP cell line was maintained in RPMI-1640 medium supplemented with 10% fetal bovine serum (FBS). The PC3 and HeLa cell lines was maintained in a 1:1 mixture of Dulbecco’s modified Eagle’s medium and F12 medium (DMEM-F12) supplemented with 10% FBS.

## Cell capture and lysis

Following standard tissue culture trypsinisation, cells were diluted to 100 cells per in PBS. Each cell line was handled separately. The initial dilution was transferred to a pre-cleaned glass slide on an inverted microscope. Between 30-50 cells were aspirated (Eppendorf Vacutip) in minimal volume and washed in 3 μL Ultrapure water (Invitrogen). The required numbers of cells were picked in minimal volume and washed again in 3 μL Ultrapure water before being transferred to a total volume of 2 μL Ultrapure water with 0.5U/ μL RNase inhibitor (Enzymantics). When 2 cell lines were mixed, the required number of cells from each line was transferred in 1.0 μL /cell line. This final volume was transferred to a lo-bind 200 μL PCR tube (Eppendorf) and snap frozen on dry ice immediately. Cells were stored at −80°C for at least 3 hr to ensure lysis.

## Reverse transcription and amplification for SS2_PCR

First strand cDNA synthesis was performed by adding 2.5 μL of a 12ng/μL stock of oligo dT barcoded primer (PCR_IVT_RTBCn or 25bpmix_BCn; IDT) to 2.5 μL of ERCC mix (6 μL 1/5000 ERCC^6^ mix 1 stock (Ambion), 85 μL nuclease free water, 5 μL RNAse inhibitor (10 U/μL Enzymatics), 4 μL 10% Triton and 100 μL dNTP mix (10 mM Roche)). A 2 μL volume of this Primer/ERCC mix was then added to the 2 μL containing the cells. This was denatured at 72°C for 3 min and immediately placed on ice. 6 μL of the first strand reaction mix, containing 0.5 μL PrimeScript reverse transcriptase enzyme mix, 2 μL 5x Primescript Buffer (both from Takara Bio), 2 μL betaine (5M Sigma), 0.25 μL DTT (100 mM Invitrogen), 0.24 μL MgCl2 (250 mM, Roche) and 0.6 μL of template switch oligo (10 μM TSO_Biotin; IDT). The reaction was kept at 25°C for 5 min, 42°C 90 min, 70°C 15 min then placed at 4°C hold. This cDNA pool was pre-amplified in 20 μL PCR reactions using 10 μL x2 HiFi Kapa PCR mix (Kapa Biosystems), 8 μL UltraPure Distilled Water (Invitrogen), 1 μL of primer PCR_F (10 μM; IDT) and 1uL of primer PCR_R (10 μM; IDT). The PCR program was 98°C 3 min, 20 cycles of 98°C 20 s, 70°C 15 s, 72°C 6 min, final incubation 72°C 5 min and hold 4°C.

## Nextera library for SS2_PCR

PCR product was adjusted to 1 ng and made into Nextera libraries as described in the Smart Seq2 protocol^15^. For the PC3/HeLacell mixture experiment amplification products were combined by sample type and used to generate a representative library.

## Lysis, reverse transcription for CS_IVT

2 μL of the Primer/ERCC mix (see above) was added to 2 μL of a first strand reagent mix (Message Amp II aRNA amplification kit: 0.6 μL First Strand buffer, 0.8 μL dNTP mix, 0.3 μL RNase Inhibitor and 0.3 μL Arrayscript). This was then added to 2 μL of cells or total RNA to make a 6 μL reaction. The RT was incubated at 42°C for 2 hr. 14 μL of second strand reagents (10.6 nuclease free water, 2 μL x10 second strand buffer, 0.8 μL dNTPs, 0.4 μL DNA polymerase and 0.2 μL RNaseH) were added and the reaction incubated at 16°C for 2 hr. cDNA was then pooled and cleaned up in accordance with the CelSeq protocol^2^.

## IVT and library finishing

IVT of CS_IVT cDNA or PCR preamplifications: At least 0.5 ng of combined template was required for a 40 μL IVT using a standard MessageAmp II kit protocol (Ambion). However, prior to purification of the aRNA 2 μl of Turbo DNase was added (Ambion). This was incubated at 37°C for 15 min, and then 39 μL of UltraPure Distilled Water (Invitrogen) was added. To fragment the aRNA 9 μL of Fragmentation solution (NEB) was added and the solution was incubated at 94°C for 90 s before addition of 10 μL of ‘stop’ solution (NEB). Fragmented aRNA was purified using a RNA Min-elute kit (Qiagen) and eluted in 15 μL of pre-warmed RNase free water (55°C). The aRNA was quantified with Qubit RNA assay (Invitrogen) and size information was obtained by Bioanalyser trace using the RNA 6000 Pico kit (Agilent).

Library finishing of aRNA by RT-PCR: 2 μl aRNA (containing approximately 1 μg) was combined with 4 μL nuclease-free water, 2 μL Primescript x5 buffer, 1 μL Second_RT primer (IDT) and 1 μL Primescript. This was incubated at 25°C for 5 min, 37°C for 30 min, 85°C for 1 min then kept at 4°C. 25 μL of Agencourt Ampure beads (Beckman Coulter) were added and incubated at 15 min at room temperature. The supernatant was then removed and replaced with PCR mix (20 μL nuclease-free water, 25μL Kapa HiFi HotStart ReadyMix (x2 Kapa Biosystems), 2.5 μL Forward primer, 2.5 μL Reverse primer. This was incubated at 98°C for 3 min followed by six cycles of: 98°C for 30 s, 65°C for 1 min, 72°C for 1 min 30 s and stored at 12°C. The resulting library was cleaned using Agencourt Ampure XP beads (Beckman Coulter) in accordance with the CelSeq protocol^2^.

## Tagmentation and library finishing

The Tn5MErev (IDT) and FC-121-1030 (IDT) oligos were resuspended to 100 μM in TE buffer. To anneal the oligos together equal volumes of 100 μM stock were combined at room temperature and mixed (final concentration of 50 μM). To load the Tn5 4 μL of the oligo mix was combined with 4 μL of the Tn5 transposase (creative-biogene, NY, USA), 2uL TPS buffer and 10 μL nuclease free water. This was incubated at 25°C for 30 min. Tagmentation of the library was achieved by combining a 20 μL solution containing 50 ng of combined PCR preamplification products with 6 μL 5x LM buffer (creative biogene) and 4 μL of the loaded Tn5. This mixture was incubated at 55°C for 20 min. 1uL of the Tn5 treated DNA was made to 10 μL with nuclease free water and combined with 6 μL PCR mix (Illumina Nextera XT kit) and 2 μL of an i7 barcoded primer (Illumina Nextera XT Index Kit) and 2 μL PCR_Lib_Primer1_i5. Note that different aliquots of this template were used to set up individual PCRs containing a different barcoded Nextera XT primer i702, i703, i704 or i705. PCR was carried out as described for Nextera kit and the product was cleaned up using Agencourt Ampure XP beads (Beckman Coulter) at vol/vol ratio of 0.6 beads to PCR product.

## Illumina high-throughput sequencing

Libraries were quantified using Qubit High Sensitivity DNA assay and size information was obtained by Bioanalyser trace using the High Sensitivity DNA assay. MiSeq runs used Illumina MiSeq Reagent Kit v3 150 Cycle kits. Read1 was a 25 bp inline barcode/Nextera Transcript Paired End 1, Read2 was the Nextera i7 8bp Barcode, Read3 was the Nextera i5 8bp Barcode and Read4 was 100bp 3’ Transcript Tag/Nextera Transcript Paired End 2.

For runs that required SS2_PCR libraries sequenced for full length sequencing we used Read1 76 bp Paired End1, Read2 Nextera i7 8 bp, Read3 Nextera i5 8 bp and Read4 76 bp Paired End 2.

## Data analysis

Data Preprocessing: Paired end Fastq files were demultiplexed using sabre (https://github.com/najoshi/sabre) via the first 8 bp of the forward read (inline barcode). Fastq files corresponding to the demultiplexed reverse read were then processed with fast x trimmer (https://github.com/agordon/fastx_toolkit) to trim 10 bp from the 5’ end (to remove any artificial sequence that may still be present from incorporation of the TSO), perform adapter trimming and a final length trim to 50 bp. We also discarded sequences with stretches of continuous Adenines greater than 25 bases in length in the 3’ half of the trimmed reads (see Supplementary File1). Mapping to the transcriptome and the ERCC reference was done using Bowtie2^16^. The transcriptome reference file was generated using a fastq file containing all known transcripts obtained from the UCSC browser^17^. We appended a 100bp stretch of Adenines to each reference sequence^9^. Mapped reads were collapsed to Gene IDs and count data was converted to transcripts per million reads (tpm) using R (http://www.R-project.org).

Mapping to the genome was also carried out using the Illumina Basespace Tophat application (https://basespace.illumina.com).

Differential expression. ERCC and PCA analysis was carried out using R and R studio (http://www.rstudio.com/). LIMMA^18^ was used to normalise the count data. Genes with less than 40 reads across samples were discarded. Pearson correlations were calculated, clustered and displayed using gplots (https://cran.rproject.org/package=gplots). We also processed the single cell data from PC3 and LNCaP cells using the SCDE analysis package^19^. SCDE was used to generate the gene specific reports in Supplementary Fig. 6 B,C. Spearman correlations were calculated, clustered and displayed using gplots.

UMI analysis. Each UMI was associated with a transcript isoform mapping identifier. Where the same UMI was associated with multiple isoforms from the same gene locus we combined the UMI frequencies.

ERCC mix1: The log2 amount of ERCC Mix 1 molecules added was plotted against the log2 absolute amount of ERCC Mix1 molecules detected with a fitted linear regression. A slope and R2 approaching one indicated that the method is detecting RNA proportional to the amount added. The estimate of the proportion of molecules detected was then estimated by plotting the mean proportion of molecules detected at each ERCC Mix 1 concentration against the absolute molecules added. A fitted linear regression with a slope very close to zero was indicative of low bias in the method. The estimate of the proportion of molecules detected by this method was then given by the intercept of a line of best fit with a slope of zero. The same method was then applied to the separated ERCC for slightly increased accuracy and to better determine patterns in outliers (data not shown).

BAM/SAM files were imported into SeqMonk v0.29.0 (http://www.bioinformatics.babraham.ac.uk/projects/seqmonk/) and aligned to 149,135 gene probes representing mRNA features in the GrCh37 genome. The raw counts for samples were normalized to the total read count and gene probe length using the SeqMonk read count quantitation tools. Normalization to total read count was done by scaling up all reads to the largest data store. Normalization to gene probe length produced reads per kilobase of transcript per million mapped reads (RPKM) for each gene probe. mRNA feature expression level reports were generated in SeqMonk and converted to CSV files for principal component analysis (PCA) using the R stats package. Only gene probes with more than 5000 reads across all single cell samples were considered for PCA. The gene expression values for the remaining 61,438 gene probes were converted to a Log base 2 scale and scaled by mean-centering and dividing by the standard deviation of each gene across all samples.

## References

1. Day, R.C., McNoe, L. & Macknight, R.C. Evaluation of global RNA amplification and its use for high-throughput transcript analysis of laser-microdissected endosperm. IntJPlant Genomics 61028 (2007).

2. Hashimshony, T., Wagner, F., Sher, N. & Yanai, I. CEL-Seq: single-cell RNA-Seq by multiplexed linear amplification. Cell Rep 2, 666-673 (2012).

3. Picelli, S. et al. Smart-seq2 for sensitive full-length transcriptome profiling in single cells. Nat Methods 10, 1096-1098 (2013).

4. Sasagawa, Y. et al. Quartz-Seq: a highly reproducible and sensitive single-cell RNA sequencing method, reveals non-genetic gene-expression heterogeneity. Genome Biol 14, R31 (2013).

5. Tang, F. et al. mRNA-Seq whole-transcriptome analysis of a single cell. Nat Methods 6, 377-382 (2009).

6. Baker, S.C. et al. The External RNA Controls Consortium: a progress report. Nat Methods 2, 731-734 (2005).

7. Islam, S. et al. Quantitative single-cell RNA-seq with unique molecular identifiers. Nat Methods 11, 163-166 (2014).

8. Marinov, G.K. et al. From single-cell to cell-pool transcriptomes: stochasticity in gene expression and RNA splicing. Genome Res 24, 496-510 (2014).

9. Pollen, A.A. et al. Low-coverage single-cell mRNA sequencing reveals cellular heterogeneity and activated signaling pathways in developing cerebral cortex. Nat Biotechnol 32, 1053-1058 (2014).

10. Ramskold, D. et al. Full-length mRNA-Seq from single-cell levels of RNA and individual circulating tumor cells. Nat Biotechnol 30, 777-782 (2012).

11. Yang, M., Loda, M. & Sytkowski, A.J. Identification of genes expressed differentially by LNCaP or PC-3 prostate cancer cell lines. Cancer Res 58, 3732-3735 (1998).

12. Grun, D., Kester, L. & van Oudenaarden, A. Validation of noise models for single-cell transcriptomics. Nat Methods 11, 637-640 (2014).

13. Zhou, M., Bhasin, A. & Reznikoff, W.S. Molecular genetic analysis of transposase-end DNA sequence recognition: cooperativity of three adjacent base-pairs in specific interaction with a mutant Tn5 transposase. J Mol Biol 276, 913-925 (1998).

14. Zechel, S., Zajac, P., Lonnerberg, P., Ibanez, C.F. & Linnarsson, S. Topographical transcriptome mapping of the mouse medial ganglionic eminence by spatially resolved RNA-seq. Genome Biol 15, 486 (2014).

15. Picelli, S. et al. Full-length RNA-seq from single cells using Smart-seq2. Nat Protoc 9, 171-181 (2014).

16. Langmead, B., Trapnell, C., Pop, M. & Salzberg, S.L. Ultrafast and memoryefficient alignment of short DNA sequences to the human genome. Genome Biol 10, R25 (2009).

17. Meyer, L.R. et al. The UCSC Genome Browser database: extensions and updates 2013. Nucleic Acids Res 41, D64-69 (2013).

18. Ritchie, M.E. et al. limma powers differential expression analyses for RNAsequencing and microarray studies. Nucleic Acids Res 43, e47 (2015).

19. Kharchenko, P.V., Silberstein, L. & Scadden, D.T. Bayesian approach to single-cell differential expression analysis. Nat Methods 11, 740-742 (2014).

